# Effects of agricultural systems on the anuran diversity in the Colombian amazon

**DOI:** 10.1101/2020.04.29.068940

**Authors:** Juan C. Diaz-Ricaurte, Nayra Camila Arriaga-Villegas, Juan David López-Coronado, Gina Ximena Macias-Garzón, Bruno F. Fiorillo

**Affiliations:** Laboratório de Ecología, Evolução e Conservação de Anfíbios e Répteis, Departamento de Ecologia, Instituto de Biociências, Universidade de São Paulo, Rua do Matão, 05508-090 São Paulo, SP, Brasil; Grupo de Investigación en Biodiversidad y Desarrollo Amazónico (BYDA), Programa de Biología, Facultad de Ciencias Básicas, Universidad de la Amazonía, Florencia, Caquetá, Colombia; Universidad de la Amazonía, Facultad de Ciencias Básicas, Programa de Biología, Florencia, Caquetá; Escola Superior de Agricultura Luiz de Queiroz, Programa de Pós Graduação em Ecologia Aplicada, Universidade de São Paulo, Piracicaba, SP, Brazil

**Keywords:** Agricultural landscapes, conservation, frogs, human-modified landscapes, habitat use

## Abstract

We provide information on the diversity of anurans from agroforestry systems in the Colombian Amazon. This area is inserted at the tropical rainforest ecosystem and consists mainly of secondary forest remnants surrounded by crops, grasslands, and agroforestry systems. From February to May 2015, we sampled anurans mainly with visual and auditory surveys. We recorded a total of 1096 individuals of 20 species of anurans from six families at the study area. The relictual forest was the richest environment, followed by Achapo and Cacao agroforestry systems. The Achapo system showed great similarity in species composition with relictual forest, however, the latter presented the highest number of exclusive species, whereas the first presented only two and Cacao system didn’t have any exclusive species. Our results show that the richness can vary between the different types of agroforestry systems and highlight their importance as management tool for anurans conservation in the Colombian Amazon.

## Introduction

Deforestation and consequent loss of biodiversity have considerably increased throughout the world in the last decades (Myers et al. 2000; Hansen et al. 2010, 2013). Such anthropic disturbances aim mostly to provide a proper structure for croplands (e.g. Lavelle et al. 1992, Mboukou-Kimbatsa et al. 1998; Barros et al. 2002). In this way, ecological studies in human-modified landscapes are extremely relevant for a major understanding of these environments as well as promoting more sustainable practices (Chazdon et al. 2009). The establishment of the agroforestry system is an alternative way for the traditional agriculture system that intents balancing agricultural development and natural landscapes’ maintenance (Izac & Sanchez 2001). Moreover, it enhances the exchange between habitat remnants (Asare et al. 2014).

Although to reduce deforestation successfully, agroforestry depends largely on local features (e.g. social and technological issues), profitability and demand of what is being produced (Angelsen & Kaimowitz, 2004), some studies have shown that shaded Cacao (*Theobroma cacao*) contributes to slow down the deforestation (Ruf & Schroth 2004). Moreover, this system usually presents a greater diversity of both plant and animal (Argôlo 2004; Schroth & Harvey 2007; Teixeira et al. 2015) than other agricultural land uses (Schroth & Harvey 2007). The same has been found for shaded coffee plantations, which can harbor a wide variety of animals, including birds, bats, and other mammals, insects, and reptiles (Somarriba et al. 2004). Despite those evidences, there is a few number of studies comparing agro-forests with other types of disturbed landscapes (Schroth & Harvey 2007).

Anurans are highly affected by habitat loss and fragmentation (mainly due to risky breeding migrations; Becker et al. 2007). In terms of land use, some systems may lead to a decrease in populations resilience and to increase extinction risks, particularly for habitat specialists (Yandi et al. 2016, Fiorillo et al. 2018). Although some species can benefit from man-made structures such as artificial water bodies, since they provide potential sites for refuge and reproduction (Brand & Snodgrass 2010; Fiorillo et al. 2019) it does not prevent that intrinsic features of disturbed habitats will affect the abundance and temporal distribution (Silva & Rossa-Feres 2011) likewise richness and composition (Wanger et al. 2010; Fiorillo et al. 2018) of anuran communities.

Piedmont region is situated located at the transition zone between the Andes mountain range and the Amazon basin, presenting high levels of biodiversity. Nevertheless, it has been the target of human settlements and thus, consequent landscape changes (Myers et al. 2000; Ricaurte et al. 2014). Considering that most amphibian species are affected in some way by habitat fragmentation and other types of disturbing, the aim of the present study was to assess the variation in anurans diversity at different agroforestry structures.

## Materials and methods

### Study area

The present study was carried out at the Amazonian Research Center Macagual - Cesar Augusto Estrada González of University of the Amazon (380 ha). It is located at 20 km from the municipality of Florencia, south of the Department of Caquetá, Colombia, (1.616667, 75.61; Datum WGS84; 300 m a.s.l; Figure 1). The mean precipitation is 3793 mm and mean annual temperature is 25.5 °C. The area is inserted at the tropical rainforest ecosystem and consists mainly of secondary forest remnants surrounded by crops, grasslands, and agroforestry systems (see below). In addition, the area remains temporarily flooded during the rainy season, a feature that partially determines the vegetation cover (Arriaga-Villegas et al. 2014). We sampled three types of agroforestry systems with different levels of complexity (vertical stratification, amount of vegetation’s cover and floristic richness and productivity (Table 1). In these sites, the floristic composition varies from timber to fruit species from the families Fabaceae, Melastomataceae, Lauraceae, Sapotaceae, Moraceae, Myristicaceae, with low presence of stubble or open areas (authors, *pers. obs*).

**Figure 1.**
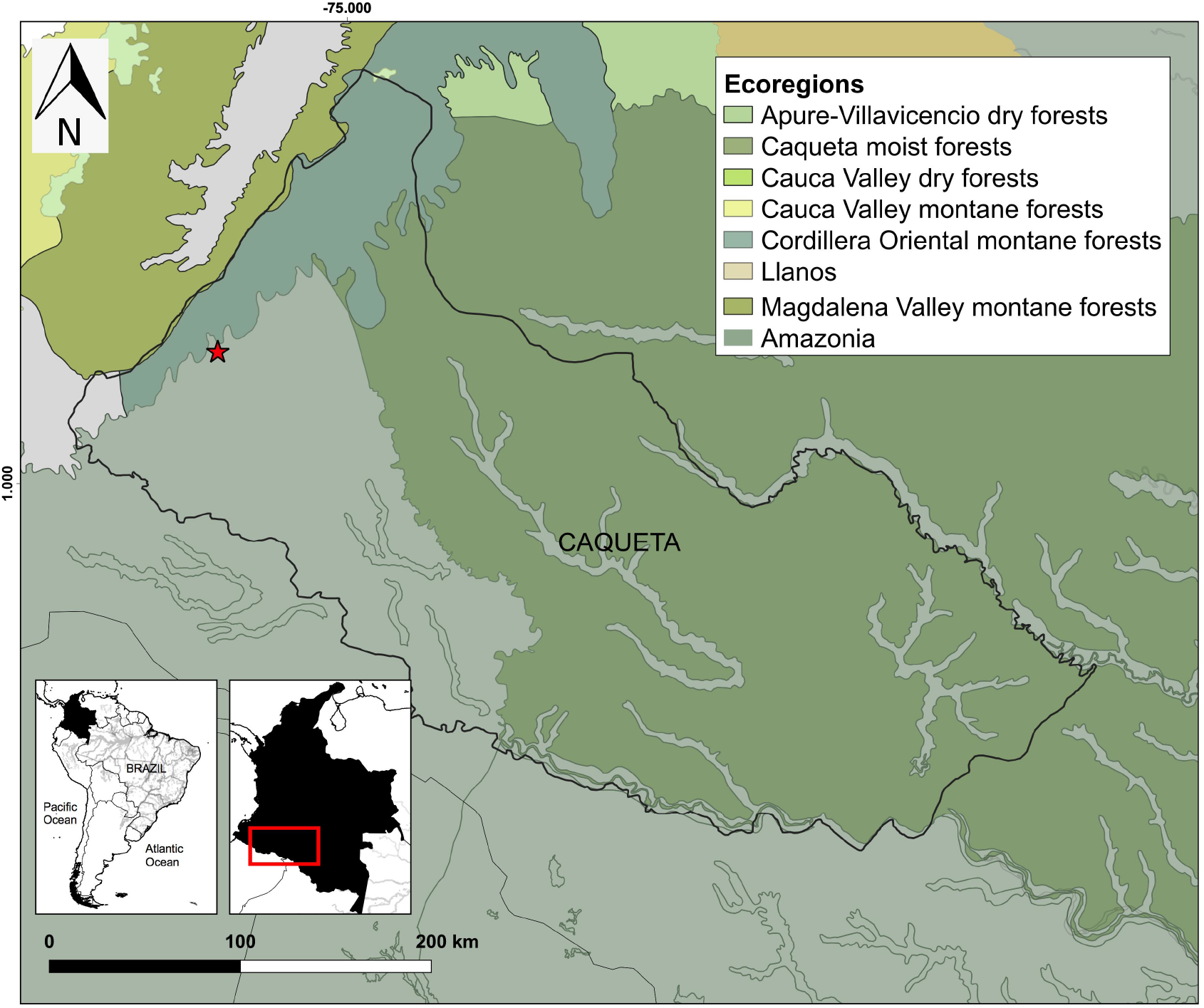
Ecoregions map of Colombia, showing the location of the Caquetá department in Colombia. Ecoregions were adapted from Dinerstein et al. (2017). Red star indicates study area location.

**Table 1.** Description of the sampled agroforestry systems in Amazon foothils, Department of Caquetá, Colombia.

| Agroforestry system | Coordinates | Elevation (m) | Vegetation |
| --- | --- | --- | --- |
| Cacao | 1.497578, -75.664791 | 235 | Agrosystem composed mostly by fruit crops, such as Azara (*Eugenia stipitata*) and Cupuassu (*Theobroma grandiflorum*), and Peach Palm (*Bactris gasipaes*). |
| Achapo | 1.500946, -75.662211 | 240 | Agroforestry system composed mainly by timber species, such as: Abarco (*Cariniana pyriformis*), Azara (*Eugenia stipitata*), Bilibil (*Guarea* sp.), Brazilian orchid tree (*Bauhinia forficata*), Common cedro (*Cedrela odorata*), Cupuassu (*Theobroma grandiflorum*). Pink poui (*Tabebuia rose*), Pink shower tree (*Cassia grandis*) and the rain tree (*Samanea saman*). |
| Relictual forest | 1.497562, -75.672103 | 243 | Forest remanent which has undergone certain transformation processes, due to deforestation for cattle breeding |

### Sampling

Data were gathered by four researchers along 8 days per month, from February to May 2015, for a total sample of 33 field trips and 26 days. At each agroforestry system, three quadrants of 70 × 20 m were randomly distributed. Within those quadrants we performed visual searches during the day (0700-1200h) and at night (19:00 - 24:00 h), totalizing a sampling effort of 99 person-hours of visual search. All individuals found were counted, identified and released.

### Data analysis

Sampling sufficiency for the different agroforestry systems was estimated through sampling coverage analysis, performed on the virtual platform iNEXT (Chao et al. 2016). This method prevents potential biases, caused by differences in species composition and/or in sampling effort of traditional species accumulation curves by random sampling. To provide the species composition and abundance distribution in all agroforestry systems we constructed a Venn diagram and abundance diagram (containing the total number of individuals observed in the field). A modification was made in the abundance diagram to include the total number of individuals captured of the species *Ameerega hahneli* (which presented a much larger number of individuals than the other species). Regarding the relative abundance of the species they were classified as abundant (more than 30 individuals found), of intermediate abundance (between 20 and 30 individuals), and rare (less than 10 individuals). To compare the species abundance from different agroforestry systems we performed a principal coordinates analysis (PCoA), using the Bray-Curtis’ similarity coefficient, and a cluster analysis, using the Simple Matching Coefficient as a measure of similarity and the Pair Group Average Method (UPGMA) as the clustering method (Sneath & Sokal 1973; Manly 1994). This analysis is useful in situations where there are more variables (species) than cases (assemblages; Sawaya et al. 2008).

## Results

A total of 1096 individuals of 20 species of anurans from six families were recorded in the study area (Table 2). Any of those is currently classified as threatened according to International Union for Conservation of Nature (IUCN), Colombian’s red list and literature (e.g. Acosta-Galvis 2000; Rueda-Almonacid et al. 2004). Most species were found in the relictual forest (RF), followed by Achapo (15 species) and Cacao (11 species) systems. Dendrobatidae was the most abundant family (500 individuals from a single species, *A. hahneli*), followed by the Leptodactylidae (252 individuals from six species) and Craugastoridae (22 individuals from two species) (Figure 2A). The Families with greatest species richness were Hylidae (seven species) and Leptodactylidae (six species), followed by Bufonidae (three species), Craugastoridae (two species), Aromobatidae and Dendrobatidae (each one with a single species; Table 2).

**Table 2.**
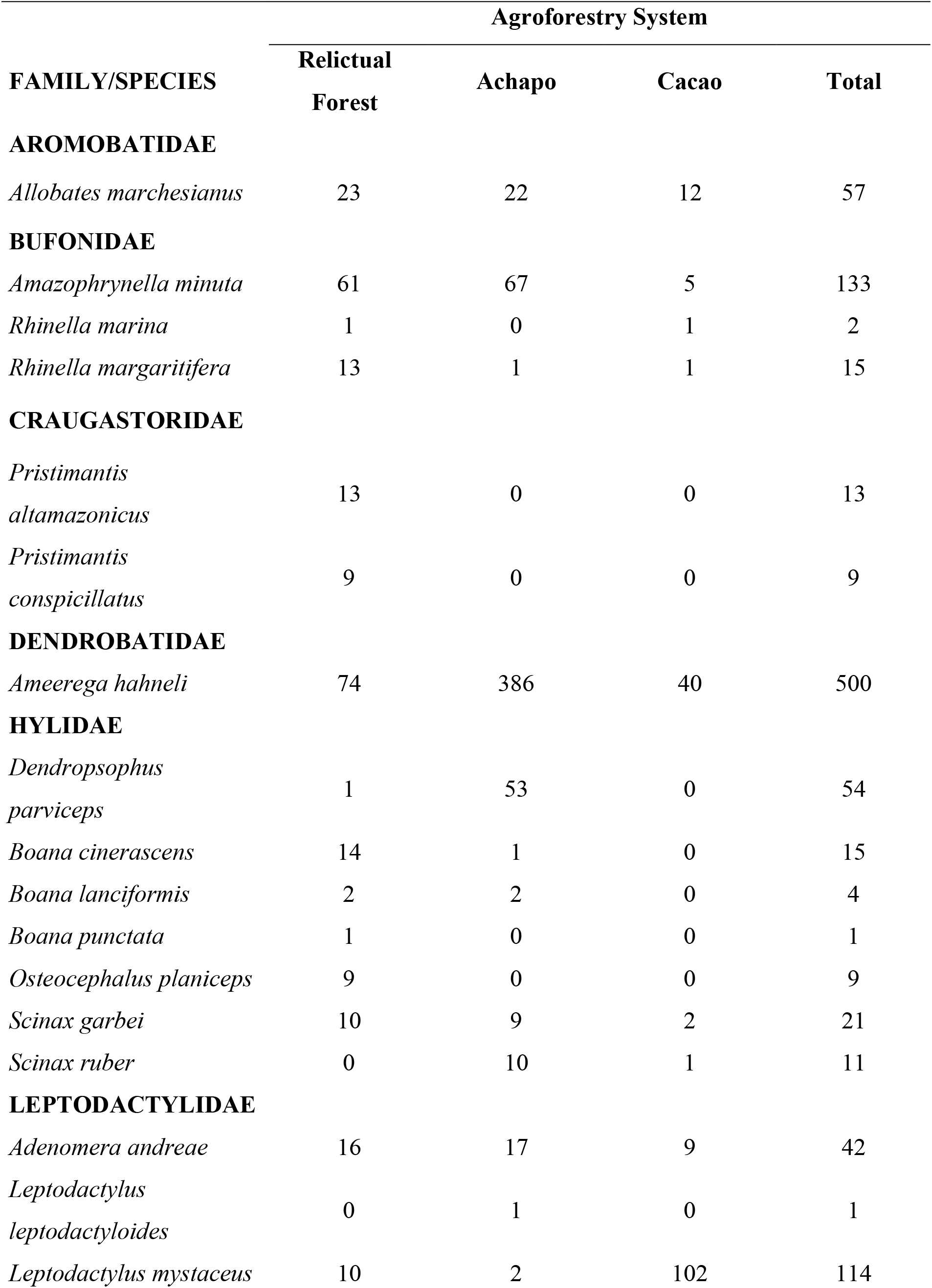
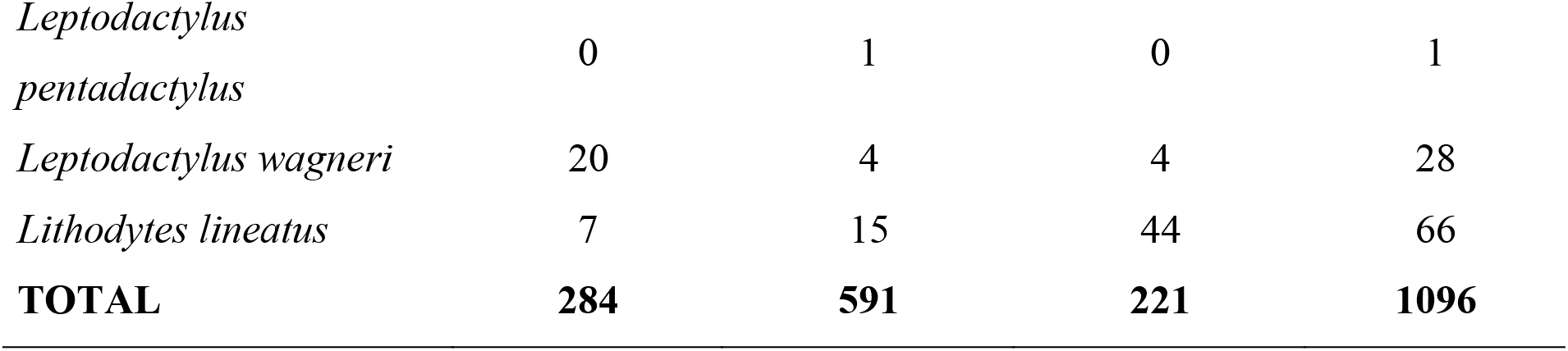
Abundance of anuran species recorded in relictual forest, Achapo and Cacao systems in the Amazonian Piedmont, Department of Caquetá, Colombia.

**Figure 2.**
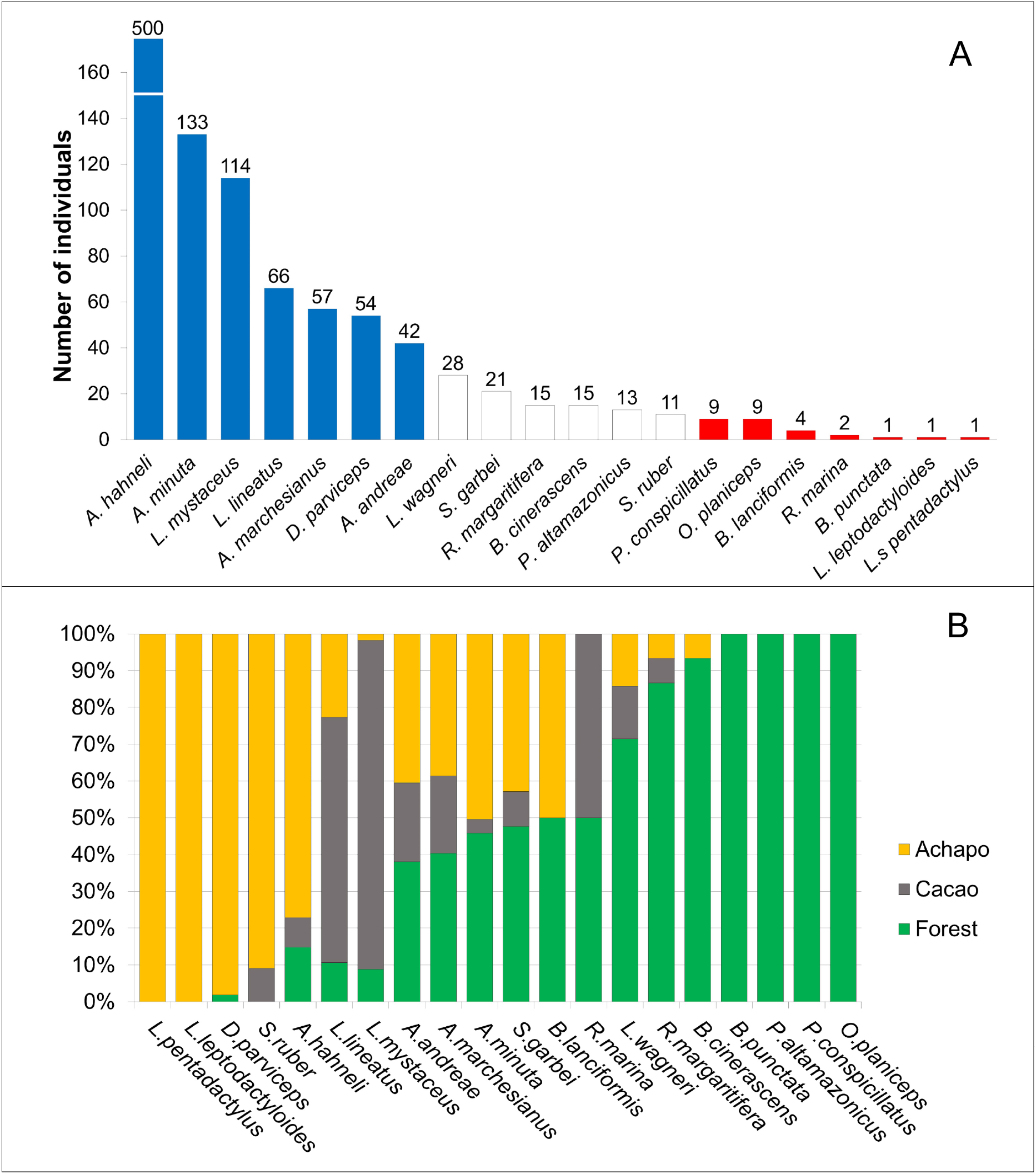
A) Distribution of abundances of anurans sampled. (absolute numbers are shown above each bar) and B) proportion of the abundance in each agroforestry system sampled at Amazonian Research Center Macagual, Department of Caquetá, Colombia.

The species richness of Achapo and RF was similar, varying from 15 to 18 species respectively. We found all species of the study area in these two environments, where *L. leptodactyloides* and *L. pentadactylus* were exclusive from Achapo, while *P. conspicillatus, P. altamazonicus, B. punctata* and *O. planiceps*, were exclusive from RF. However, despite of most species had been found at both, they presented great contrast in terms of abundance (Figure 2B). The species *D. parviceps* was almost exclusively found in Achapo (more than 98% of individuals) showing high affinity for this type of agroforestry system. Only nine species shared all the three systems, whereas three species were common in RF and Achapo (*D. parviceps, B. cinerascens, B. lanciformis*), one in Achapo and Cacao (*S. ruber*) and one in Relictual forest and Cacao (*R. marina*). The system with the highest number of exclusive species was the RF (*P. altamazonicus, P. conspicillatus, O. planiceps, B. punctata*), whereas only two species were exclusive of Achapo (*L. leptodactyloides* and *L. pentadactylus*) and none was exclusive in Cacao.

The sampling coverage analysis indicates that our sampling effort was enough to sample nearly all species that occur in the region, as well as in each system (Figure 3). Sampling in the different systems reached coverage of 0.97 in the forest, 0.97 in the Cacao and 0.99 in the Achapo (Figure 4).

**Figure 3.**
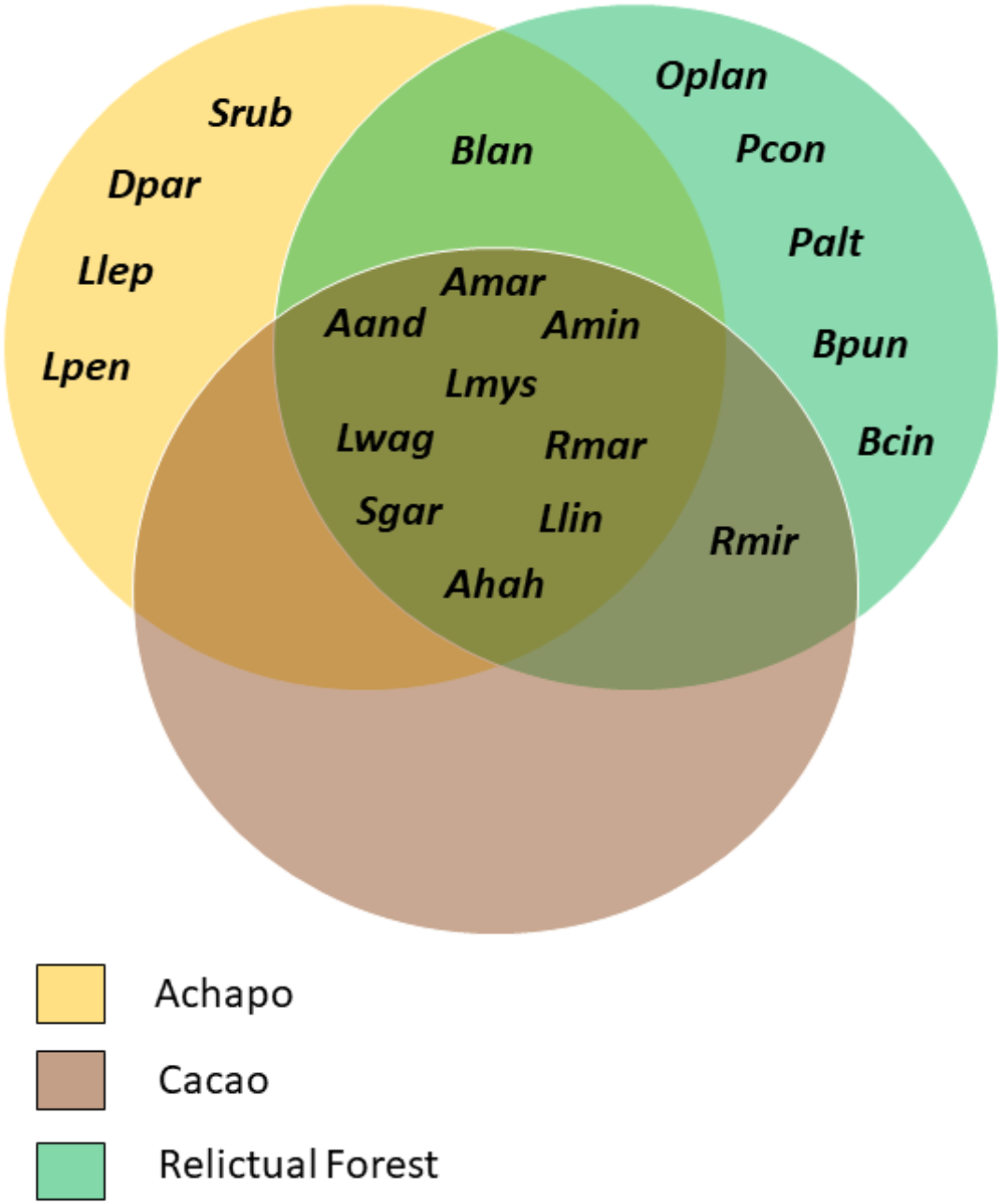
Venn diagram of the intersections between the composition of each of the three agroforestry systems sampled (Achapo, Cacao, and relictual forests). Aand: *Adenomera andreae*, Ahah: *Ameerega hahneli*, Amar: *Allobates marchesianus*, Amin: *Amazophrynella minuta*, Bcin: *Boana cinerascens*, Blan: *Boana lanciformis*, Blan: *Boana punctata*, Dpar: *Dendropsophus parviceps*, Llep: *Leptodactylus leptodactyloides*, Lmys: *Leptodactylus mystaceus*, Lpen: *Leptodactylus pentadactylus*, Lwag: *Leptodactylus wagneri*, Llin: *Lithodytes lineatus*, Oplan: *Osteocephalus planiceps*, Palt: *Pristimantis altamazonicus*, Pcon: *Pristimantis conspicillatus*, Sgar: *Scinax garbei*, Srub: *Scinax ruber*, Rmar: *Rhinella margaritifera*, Rmri: *Rhinella marina*. Obs: when more than 90% of given species was found in a single system (e.g. *Boana cinerascens*), it was considered as exclusive in the diagram.

**Figure 4.**
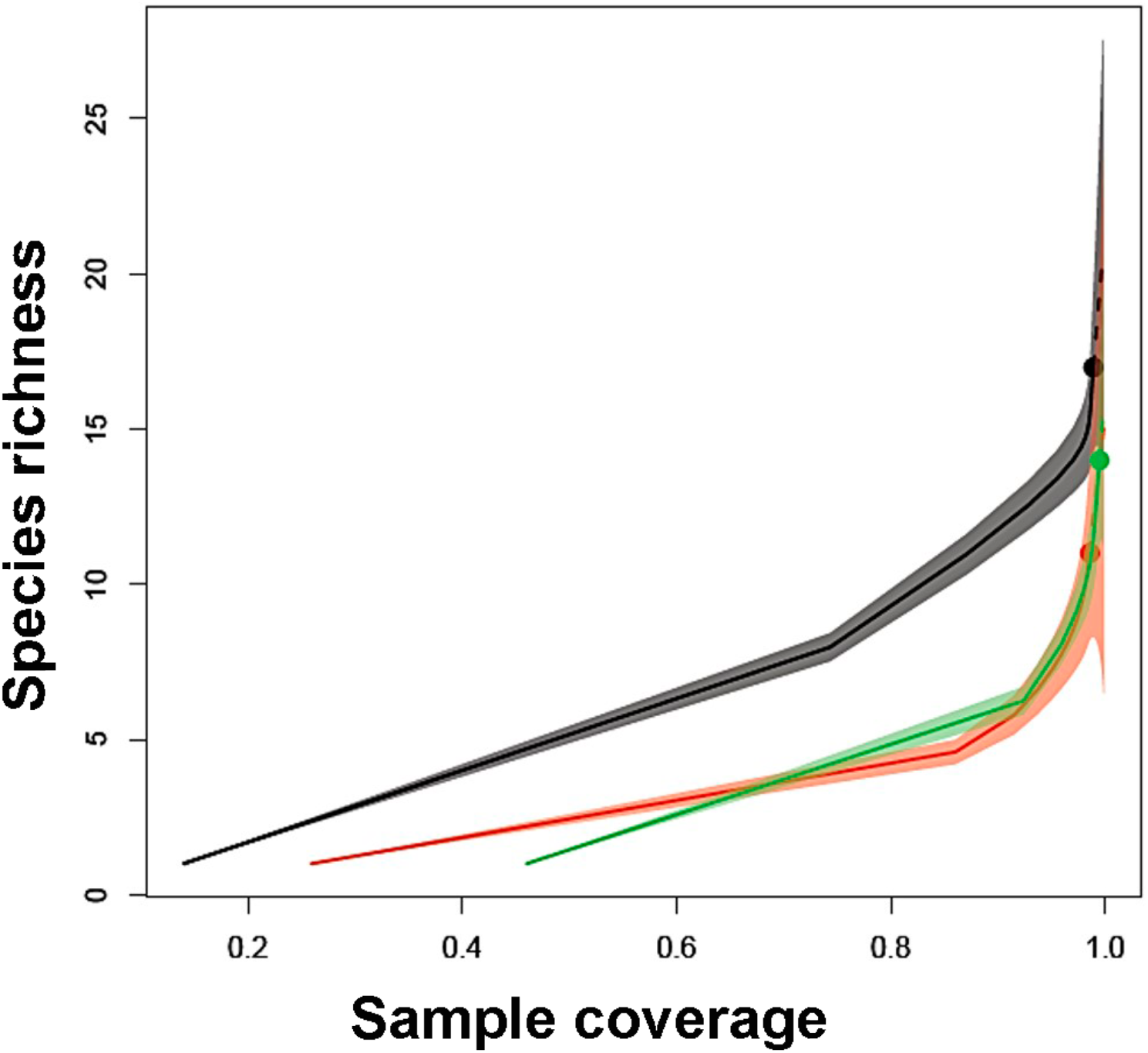
Sampling coverage curves for each agroforestry systems. Black: Relictual forest, Red: Cacao, Green: Achapo.

The first two axes of the Principal Coordinate Analysis (PCoA; Figure 5) together explained 46.95 % of the total data variance (axis 1: eigenvalue = 1.69 and 23.36 % of variance; axis 2: eigenvalue = 1.51 and 23.59 % of variance). The PCoA axis 1 ordinated species associated with the RF from those associated to Achapo, where generalist species (including those found at the Cacao system) remained at the center. The PCoA axis 2 ordinated rare and abundant species (Figure 5).

**Figure 5.**
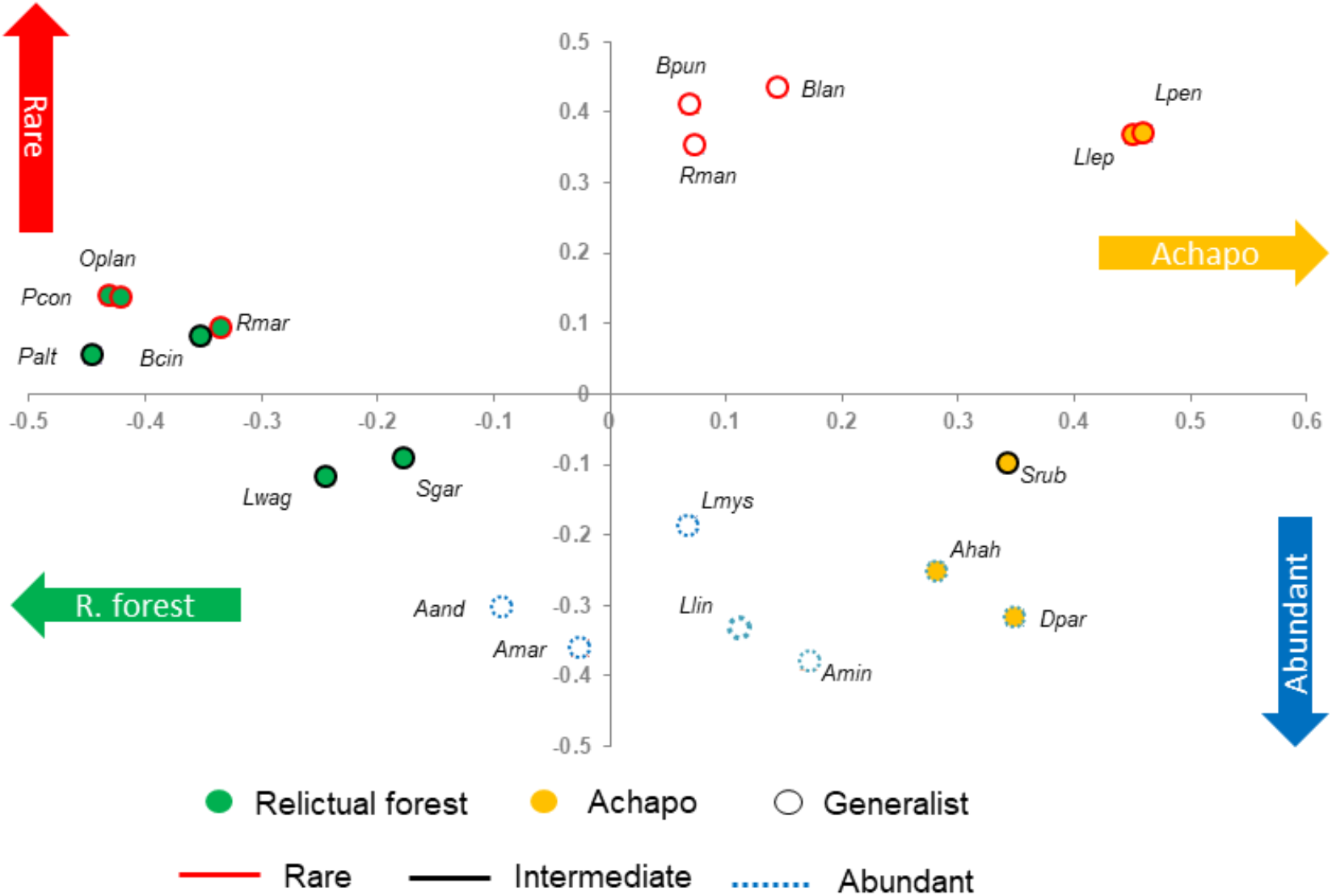
Principal coordinate analysis (PCoA) of anuran abundance, for each species recorded at Amazonian Research Center Macagual, Department of Caquetá, Colombia.

## Discussion

We found a higher species richness of anurans in RF, followed by Achapo and Cacao, the former with the largest number of exclusive species. The Achapo system is composed by timber and fruit species and presents small flooded areas which might provide a variety of microhabitats, food, shelter and potential breeding sites for anurans and other animals. Furthermore, local climatic conditions are known to be correlated with anurans diversity, what can be related to their eco-physiological constraints (Valdujo et al. 2013). Some studies have shown that the shading and variety of soil types in agroforestry systems affect local climatic conditions (Lin 2007). This suggests that features related to system structure, such as vertical stratification can indirectly affect anurans diversity.

The Cacao system was the wettest site and presented the greatest degree of anthropic disturbance (near built areas). According to that, the most abundant species in this environment was *L. mystaceus* which is considered a generalist species, known by its great capacity to adapt to altered habitats (Heyer & Bellin 1973; IUCN 2015). Although *S. ruber* is relatively rare (N = 11) in the community, it was more abundant at the Cacao System either. This species is known to occur from open to forested environments and even urban-rural areas (Duellman 1978; Duellman & Wiens 1993).

*Ameerega hahneli* was the most abundant species in the three agroforestry systems. This species is primarily terrestrial and inhabits primary and secondary forests. It uses mainly palm branches and fallen leaves as microhabitats (Carvalho 2011). The Achapo system was where it the largest number of individuals of this species was found, what may be due to this system provide a thick leaf-litter layer, favoring species foraging behaviour and reproduction (Rodríguez & Duellman 1994; Haddad & Martins 1994; Duellman 2005; Lötters and Mutschmann 2007; Vásquez et al. 2013). The same seems to be the case of the bufonid species *A. minuta* (Miranda et al. 2015; Rojas et al. 2015). The Craugastoridae family showed the lowest abundance and richness (only 22 individuals of two species at the community, both exclusive from RF system, see Table 1). This is probably related to the low capacity of some species to adapt in disturbed environments (Guo 2000; Lips et al. 2005). The Hylidae family was the richest one at the study area (seven species). This family is considered one of the most diverse among the anurans (Frost 2020) and presents a great number of reproductive modes associated from the vegetation to floor, reflecting adaptations toward habitat occupation (e.g. Haddad & Prado 2005). In contrast, Leptodactylids (second most diverse family of the study area with six species), are frequently associated with the floor, generally lacking arboreal adaptations (Haddad & Prado 2005). Species of this family were usually associated to exposed water bodies (temporary or permanent). These microhabitats are often used by species of this family for reproduction and are common at the three environments of the study area. These results show how anurans life history from Department of Caquetá can be determinant regarding their colonization at altered environments (Rodriguez & Duellman 1994).

Although no threatened species has been found at the study area, the replacement of natural forests by agricultural landscapes (e.g. pastures, palm plantations; (Etter et al. 2006; Garcia-Ulloa et al. 2012), contributes to both extinguish forest specialists (such as *O. planiceps*, *P. altamazonicus*, *P. conspicillatus*, *B. punctata*, and *B. cinerascens*, found exclusively in relictual forests) as well as to facilitate the occupation by generalist ones (Marvier et al. 2004). Forest habitats present a great structural complexity (arboreal and understory remains) which provides a variety of microhabitats (Urbina & Pérez 2002; Jose 2012; Palacios et al. 2013). Both Achapo and Cacao seem to work as transition environments, which arise from forest fragmentation. Fiorillo et al. (2018) found that anurans diversity from peach palm plantations and secondary forests that surrounded it was very similar. On the other hand a site of banana plantation from the same study presented lesser richness than both environments (peach palm plantations and secondary forests) and none exclusive species. These results are very similar to that showed at the present study and indicate that anurans diversity of disturbed habitats could be related to the interplay between populations source (natural landscapes) and intrinsic traits of disturbed habitats (e.g. crops, agroforestry systems).

We conclude that the agroforestry systems can be important management tools for anurans conservation. Their multi-stratified structures are able to provide shelter and suitable reproductive conditions for a great number of species. However, it is important to have in mind that, some of those systems can harbor richest communities than others. Additionally, the effectiveness of biological conservation depends largely on the agricultural matrix and how it is managed (Perfecto & Vandermeer 2008; Jose 2009). The assessment of landscape changing on biodiversity and species natural history can be critical for promoting more sustainable practices.

## Acknowledgements

We thank University of Amazon for allowing our fieldwork at Amazonian Research Center Macagual - Cesar Augusto Estrada González. We thanks to Maykol Olarte, Ricardo Rodriguez, Rolland D. Díaz-Morales and Karen Garcia for assistance in field work. This study was financed in part by the Coordenação de Aperfeiçoamento de Pessoal de Nível Superior – Brasil (CAPES) – Finance Code 001. Finally, we thank Maria Juana Parada Sierra for the English revision.

## Disclosure statement

No potential conflict of interest was reported by the authors.

## Author contributions

JCDR planned the study; JCDR, NCAV and JDLC collected the data; GXMG financed fieldwork; JCDR and BFF analyzed the data; JCDR led the writing of the manuscript. JCDR wrote the original draft; JCDR and BFF contributed critically to the manuscript. All authors approved the final version of this manuscript for publication.

## Notes

### Competing Interest Statement

The authors have declared no competing interest.

